# Sashimi plots: Quantitative visualization of alternative isoform expression from RNA-seq data

**DOI:** 10.1101/002576

**Authors:** Yarden Katz, Eric T. Wang, Jacob Silterra, Schraga Schwartz, Bang Wong, Helga Thorvaldsdóttir, James T. Robinson, Jill P. Mesirov, Edoardo M. Airoldi, Christopher B. Burge

**Author notes:** These authors contributed equally (Yarden Katz, Eric T. Wang, Jacob Silterra).

## To the Editor

Analysis of RNA sequencing (RNA-Seq) data revealed that the vast majority of human genes express multiple mRNA isoforms, produced by alternative pre-mRNA splicing and other mechanisms, and that most alternative isoforms vary in expression between human tissues (Pan et al., 2008; Wang et al., 2008). As RNA-Seq datasets grow in size, it remains challenging to visualize isoform expression across multiple samples. We present Sashimi plots, a quantitative multi-sample visualization of RNA-Seq reads aligned to gene annotations, which enables quantitative comparison of isoform usage across samples or experimental conditions. Given an input annotation and spliced alignments of reads from a sample, a region of interest is visualized in a Sashimi plot as follows: (i) alignments in exons are represented as read densities (optionally normalized by length of genomic region and coverage), and (ii) splice junction reads are drawn as arcs connecting a pair of exons, where arc width is drawn proportional to the number of reads aligning to the junction (or to the log of this number) (Figure 1).

**Figure 1:**
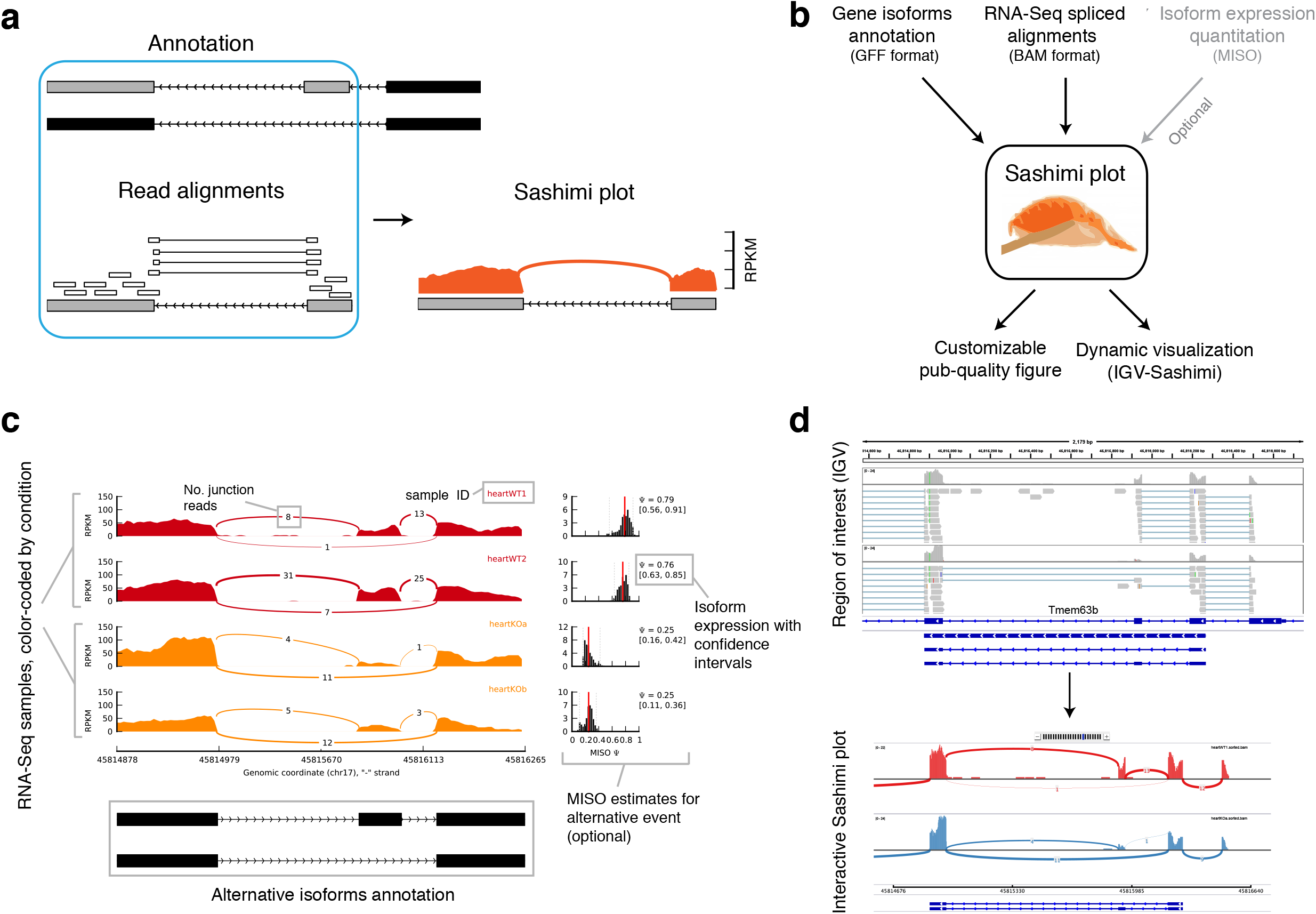
**(a)** Anatomy of a Sashimi plot. Gene model annotation containing two isoforms differing by inclusion/exclusion of middle exon. Sashimi plot for the two grey exons (blue boxed region) is shown, where genomic reads are converted into read densities (per-base expression as y-axis value) and junction reads are plotted as arcs whose width is proportional to the number of reads aligned to the junction spanning the exons connected by arc. **(b)** Inputs required for making a Sashimi plot. Gene model annotations (in GFF format), RNA-Seq read alignments (BAM format) and optionally isoform expression estimates (by MISO) are used to make Sashimi plots. Sashimi plots can be made with a stand-alone program that makes customizable publication quality figures, or dynamically from the IGV browser. **(c)** Sashimi plot (stand-alone) for alternatively spliced exon and flanking exons in four samples (colored by experimental condition). Right: optional isoform expression information produced by MISO. **(d)** Genomic region of interest in IGV along with two alignment tracks (top) from which a Sashimi plot is generated on the fly (bottom). Resulting Sashimi plot scales/resolution are set interactively by the user.

Sashimi plots require as input spliced alignments (stored in the SAM/BAM format) and gene model annotations (in GFF format (Stein, 2010)), obtainable from databases such as Ensembl or custom-made by the user (Figure 1b). Two implementations of Sashimi plots are available: (1) a stand-alone command line implementation for producing customizable publication-quality figures, and (2) an implementation built into the Integrated Genome Viewer (IGV) browser (Thorvalds
dóttir et al., 2013), IGV-Sashimi, which enables dynamic creation of Sashimi plots for any genomic region of interest, suitable for exploratory analysis of isoform usage across experiments (Figure 1b). Isoform expression estimates generated by the MISO algorithm (Katz et al., 2010) are optionally plotted in Sashimi plots.

A Sashimi plot generated by the stand-alone program for four RNA-Seq samples is shown in Figure 1c. Samples are color-coded by condition, with RNA-Seq samples from wild type mice in red and mouse heart tissues depleted for the splicing factor Muscleblind1 (‘heartKOa’, ‘heartKOb’) in orange. Read densities across exons are quantified in RPKM units (Mortazavi et al., 2008) and junction reads are plotted as arcs that are annotated with the raw number of junction reads present in each sample. Alternative isoforms from the input annotation are shown at bottom. The plot highlights the differential splicing of the middle exon, which appears to be predominantly included in the wild type samples but mostly excluded in the knockout samples. This difference is confirmed by the MISO estimates for the inclusion of the exon (Figure 1c), which indicate that inclusion levels for the exon (quantified as ‘Percent Spliced In’ or Ψ, as in (Katz et al., 2010)) is ∼77% in wild type samples and only ∼25% in the knockout samples. Users can customize the scales, colours, labels and other features of the plot through a text settings file.

An IGV-Sashimi plot for the genomic region containing the same alternative exon is shown in Figure 1d, with one wild type heart sample shown in red and one knockout heart sample in blue. The GFF annotation of the alternatively spliced exon is shown in the lower panel, and RefSeq canonical transcripts for the gene are shown above. The boundaries of the Sashimi plot are determined by the region of interest shown in the IGV browser window, and can be altered to include more or fewer exons using the zoom in/out feature of the browser. As in Figure 1c, the raw junction read counts are shown on top of each junction arc.

The Sashimi plot code base is free and open-source (available via GitHub), and can be used to combine isoform expression levels with other genomic data. The Sashimi plot code base was recently adapted to display splicing quantitative trait loci (‘sQTL’) alongside genotypic information (Wu et al., 2014).

Sashimi plots can aid in visualization of alternative splicing for use in figures, or for rapid surveying of genomic regions for differential isoform usage across multiple samples. Expression information and other genomic datasets can be integrated into Sashimi plots either programmatically or as tracks through the IGV browser.

## Software documentation and download information

The Sashimi plot software and documentation main page:

http://genes.mit.edu/burgelab/miso/docs/sashimi.html

IGV-Sashimi section from IGV browser documentation:

http://www.broadinstitute.org/software/igv/Sashimi

Source code for stand-alone Sashimi software and IGV-Sashimi is available at the following GitHub repositories:

http://github.com/yarden/MISO

http://github.com/broadinstitute/IGV

## Acknowledgements

We thank V. Butty and N. Robine for insightful discussions.

## References

Katz, Y., Wang, E.T., Airoldi, E.M., and Burge, C.B. (2010). Analysis and design of RNA sequencing experiments for identifying isoform regulation. Nature Methods 7, 1009–1015.

Mortazavi, A., Williams, B.A., McCue, K., Schaeffer, L., and Wold, B. (2008). Mapping and quantifying mammalian transcriptomes by RNA-Seq. Nature methods 5, 621–628.

Pan, Q., Shai, O., Lee, L.J., Frey, B.J., and Blencowe, B.J. (2008). Deep surveying of alternative splicing complexity in the human transcriptome by high-throughput sequencing. Nature genetics 40, 1413–1415.

Stein, L. (2010). Generic Feature Format, Version 3. Sequence Ontology Project, 1–18.

Thorvaldsdóttir, H., Robinson, J.T., and Mesirov, J.P. (2013). Integrative Genomics Viewer (IGV): high-performance genomics data visualization and exploration. Briefings in bioinformatics 14, 178–192.

Wang, E.T., Sandberg, R., Luo, S., Khrebtukova, I., Zhang, L., Mayr, C., Kingsmore, S.F., Schroth, G.P., and Burge, C.B. (2008). Alternative isoform regulation in human tissue transcriptomes. Nature 456, 470–476.

Wu, E., Nance, T., and Montgomery, S.B. (2014). SplicePlot: a utility for visualizing splicing quantitative trait loci. Bioinformatics (Oxford, England).

